# Nucleotide-decoupled G proteins reveal the role of G protein conformation in receptor-G protein selectivity

**DOI:** 10.1101/2022.05.25.493498

**Authors:** Wonjo Jang, Sumin Lu, Nevin A. Lambert

**Author notes:** Co-corresponding authors: Wonjo Jang, Department of Pharmacology and Toxicology, Medical College of Georgia, Augusta University, Augusta, Georgia, 30912 USA, Nevin Lambert, Department of Pharmacology and Toxicology, Medical College of Georgia, Augusta University, Augusta, Georgia, 30912 USA.

## Abstract

G protein-coupled receptors (GPCRs) selectively activate at least one of the four families of heterotrimeric G proteins to transduce environmental cues, but the mechanistic basis of coupling selectivity remains unclear. Structural studies have emphasized structural complementarity of GPCR complexes with nucleotide-free G proteins, but it has also been suggested that selectivity may be determined by intermediate activation processes that occur prior to nucleotide release. To test these ideas we have studied coupling to nucleotide- decoupled G protein variants, which can adopt conformations similar to receptor-bound G proteins without the need for nucleotide release. We find that selectivity is significantly degraded when nucleotide release is not required for GPCR-G protein complex formation, to the extent that most GPCRs interact with most nucleotide-decoupled G proteins. These findings demonstrate the absence of absolute structural incompatibility between most GPCRs and G proteins, and are consistent with the hypothesis that high-energy intermediate state complexes are involved in coupling selectivity.

## Introduction

G protein-coupled receptors (GPCRs) sense extracellular cues and mediate many of their physiological responses by activating heterotrimeric G proteins^1,2^. Agonist-bound GPCRs adopt active state conformations and associate with GDP-bound heterotrimeric G proteins, which facilitates release of GDP from the Gα subunit. This is followed by binding of GTP and subsequent dissociation of the heterotrimer. Despite the vast number and diversity of external stimuli GPCRs share only four families of G proteins. Because members of each family interact with different intracellular effectors to produce distinct physiological responses, GPCRs must selectively activate specific subsets of G protein heterotrimers.

The mechanistic basis of coupling selectivity has been studied extensively using receptor and G protein chimeras and site-directed mutagenesis^3,4^. More recently, structural characterization of GPCR-G protein complexes has identified elements that are important for activation of specific G proteins^5-8^. However, structural studies require stable agonist-receptor-G protein ternary complexes, and this almost always requires a nucleotide-free G protein. Therefore, such studies are largely limited to a state of the receptor-G protein complex that exists *after* nucleotide release has already occurred, in other words after the critical step in G protein activation. Receptor-mediated nucleotide release involves extensive conformational rearrangement of the Gα subunit that is unlikely to take place in a single step^9^, and it has been suggested that G protein selection occurs at an earlier point involving an “intermediate state” GPCR-G protein complex that is too energetically unstable to be captured by standard structural approaches^10,11^. A few structural and biophysical studies have begun to characterize possible intermediate state complexes^10,12^, but evidence that such complexes are involved in coupling selectivity is lacking. Here we approach this question using G protein mutants where binding to receptors is decoupled from nucleotide release, and many of the structural changes associated with G protein activation are present prior to association with a receptor^13^. We find that receptor-G protein interaction selectivity is significantly degraded with such mutants. Moreover, even in the absence of activating ligands, most receptors spontaneously form complexes with cognate nucleotide-decoupled G proteins, implying that significant basal sampling of active conformations is a near-universal feature of GPCRs. These findings suggest that nucleotide- decoupled G proteins can bypass intermediate steps that normally play a role in coupling selectivity, lending support to the idea that selectivity arises, at least in part, from transient intermediate state receptor-G protein complexes.

## Results

*Receptor association and nucleotide exchange are decoupled in 4A mutant G proteins*. Biophysical and structural studies of GPCR-G protein complexes have revealed that Gα subunits undergo large scale conformational changes that lead to GDP release^7,14^. The C-terminal helix 5 (H5) of Gα interacts with the transmembrane core of active receptors, requiring a rigid-body translation of this helix towards the receptor and rotation by 60 degrees^7^. This engagement, likely in concert with receptor interactions with the Gα N terminus, triggers several conformational changes that promote GDP release. These include: (1) disruption of a hydrophobic core comprised of H5, H1, and the S1-S3 strands; (2) increased disorder of the s6h5 loop (TCAT motif), which directly contacts the guanine ring of GDP; and (3) increased disorder of the P loop, which coordinates the β-phosphate of GDP^9^. In addition, separation of the Ras-homology and α-helical domains is necessary but not sufficient for GDP release^14^. These changes in Gα are energetically costly^15^, and presumably are compensated by formation of new contacts with the active receptor.

To test the idea that these changes might play a role in coupling selectivity we studied interactions between receptors and G protein mutants that mimicked the receptor-bound state. For this purpose, we chose a previously characterized mutant design^13^ wherein the H5 helix is extended by inserting four alanine residues (4A) between H5.5 and H5.6 (according to the <u>C</u>ommon <u>G</u>α <u>N</u>umbering (CGN) system)^16^. This insertion disrupts the H5-H1-S1-3 hydrophobic core and adds one helical turn, lengthening H5. The conserved residue F^H5.8^ is extracted from the hydrophobic core of the Gα_i1_ 4A mutant, as is the case in nucleotide-free receptor-bound structures (Fig. 1). Importantly, the G_i1_ 4A mutant remained bound to active rhodopsin even in the presence of guanine nucleotides^13^, suggesting that this modification decouples nucleotide binding from receptor association.

**Figure 1.**
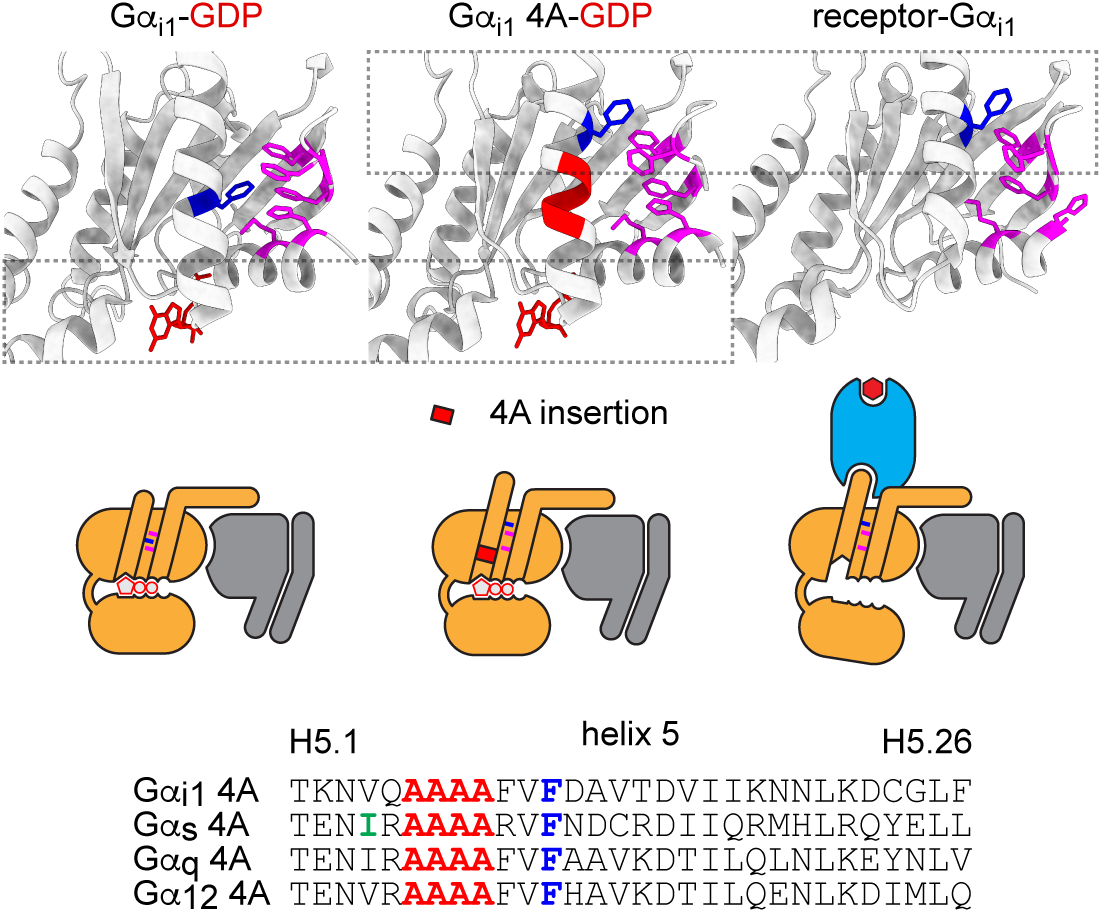
Structural features of nucleotide-decoupled G proteins. GDP-bound Gα subunits (*top left*; PDB 1GP2) possess a hydrophobic core comprised of side chains of residues in alpha helix 1 (H1), and beta strands 1-3 (S1-3; *magenta*), as well as residues in helix 5 (H5), including F^H5.8^ (*dark blue*). In structures of nucleotide-free Gα_i1_ subunits bound to active receptors (*top right*; PDB 7CMV) this hydrophobic core is disrupted, and F^H5.8^ is extracted. G protein 4A mutants (*top center*; PDB 5KDO) include four alanine residues (*red*) between H5.5 and H5.6 which extends H5, disrupts the hydrophobic core, and extracts F^H5.8^. Receptor-binding regions of 4A mutants are similar to those of receptor-bound Gα subunits, whereas GDP-binding regions of 4A mutants are similar to those of GDP-bound wild-type Gα subunits (*dashed rectangles*). At *bottom* are primary sequences of H5 in 4A mutants of Gα_i1_, Gα_s_, Gα_q_ and Gα_12_. Highlighted in *green* is residue I^H5.4^ in Gα_s_, which is mutated to alanine in mini Gs.

We first studied interactions of wild-type (wt) G_i1_ and G_i1_ 4A with a prototypical ligand-activated GPCR, the M_4_ muscarinic acetylcholine receptor (M_4_R). We measured bioluminescence resonance energy transfer (BRET) between M_4_R fused to *Renilla* luciferase (Rluc8) and Gβ_1_ and Gγ_2_ subunits fused to complementary fragments of Venus fluorescent protein, thus leaving Gα subunits untagged (Fig. 2A). To minimize background from endogenous G proteins we used HEK 293 cells in which G_s/olf_, G_q/11_ and G_12/13_ family G proteins had been deleted using CRISPR/Cas9-mediated gene editing (3GKO)^17^. Cells were permeabilized with digitonin and either treated with apyrase to remove endogenous nucleotides or supplemented with exogenous nucleotides. In the absence of nucleotides acetylcholine (Ach) elicited a large BRET increase between M_4_R-Rluc8 and wt G_i1_, which was quickly reversed upon subsequent addition of the apyrase-resistant GDP analog GDPβS (Fig. 2A). In contrast, Ach-induced BRET between M_4_R-Rluc8 and G_i1_ 4A was largely resistant to GDPβS (Fig. 2A). The absolute magnitude of agonist-induced BRET was smaller for G_i1_ 4A than wt G_i1_ because the G_i1_ 4A signal started from a higher baseline (see below). Similar results were obtained with the apyrase-resistant GTP analog GTPγS, except that agonist-induced BRET between M_4_R-Rluc8 and G_i1_ 4A was only partially resistant to GTPγS (Fig. S1). We suspected that GTPγS decreased BRET between M_4_R-Rluc8 and G_i1_ 4A by promoting dissociation of Venus-Gβγ from Gα_i1_ 4A. Consistent with this notion, BRET between M_4_R-Rluc8 and G_i1_ 4A bearing an additional mutation (R208S) to prevent GTP-induced heterotrimer dissociation was largely resistant to GTPγS (Fig. S1)^18^. These results are consistent with how G_i1_ 4A interacts with light-activated rhodopsin^13^, and show that the G_i1_ 4A interaction with the ligand-activated M_4_R-Rluc8 is also decoupled from nucleotide binding.

**Figure 2.**
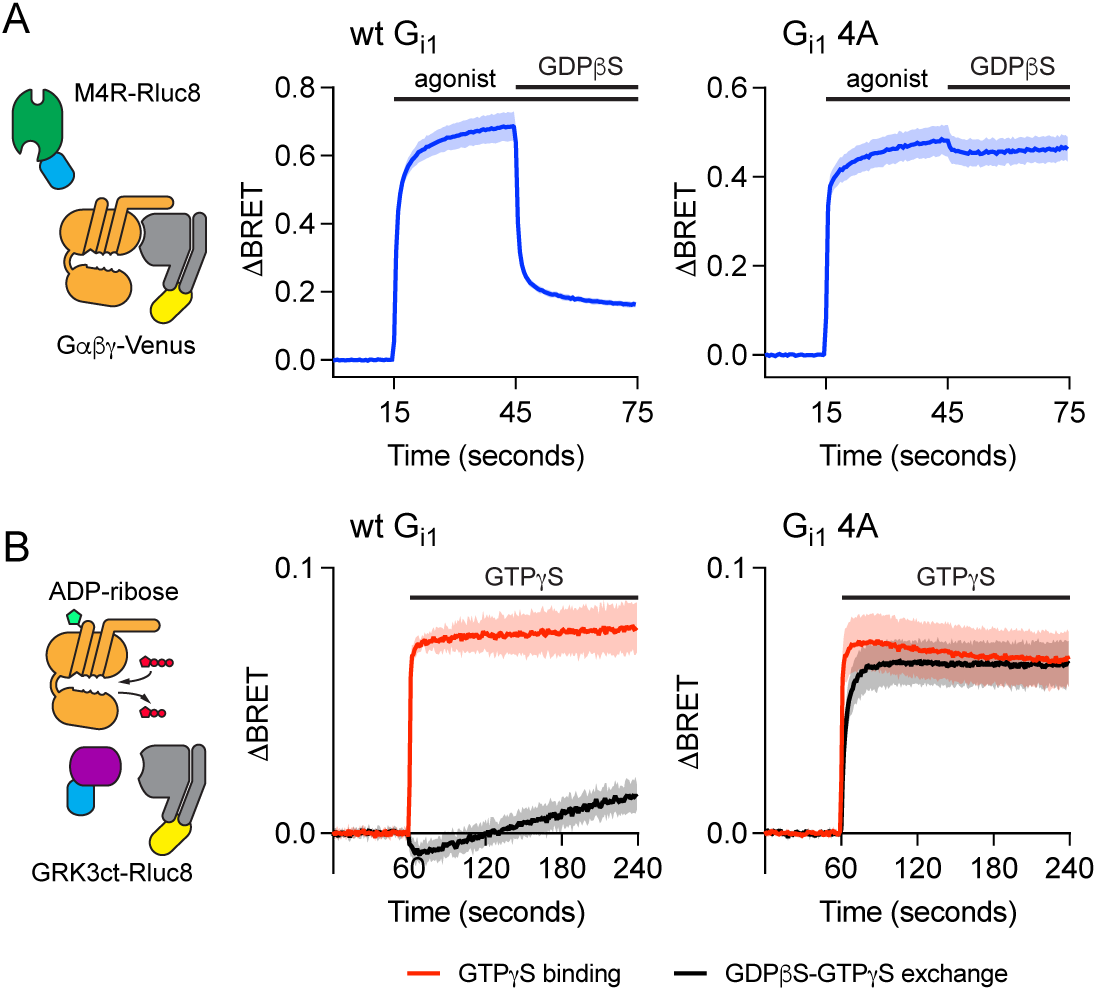
Functional properties of nucleotide-decoupled G proteins. *A*) BRET between M_4_R- Rluc8 and wild-type (wt) G_i1_ heterotrimers induced by the agonist acetylcholine (100 µM) in the absence of nucleotides is largely reversed by addition of GDPβS (10 µM), whereas BRET between M_4_R-Rluc8 and G_i1_ 4A heterotrimers is largely resistant to GDPβS (mean ± 95% C.I.; *n*=12). *B*) Wild-type G_i1_ exchanges GDPβS for GTPγS slowly, whereas G_i1_ 4A exchanges nucleotides more rapidly. The rate of GTPγS binding to nucleotide-free heterotrimers is similar for wt and 4A mutant G_i1_. Gα_i1_ subunits were ADP-ribosylated by pertussis toxin to prevent receptor-catalyzed nucleotide release; GTPγS binding was monitored indirectly using a membrane-bound free Gβγ sensor GRKct-Rluc8 after incubation in apyrase alone (for GTPγS binding) or apyrase plus GDBβS (for GDPβS-GTPγS exchange; mean ± 95% C.I.; *n*=7-14).

Isolated Gα_i1_ 4A subunits have been shown to spontaneously exchange nucleotides more rapidly than wt Gα_i1_^13^. We next asked if Gα_i1_ 4A subunits also exchange nucleotides rapidly when complexed with Gβγ dimers in cell membranes. For this purpose, we used a protocol where GDPβS-GTPγS exchange was monitored indirectly with a membrane-anchored free Gβγ sensor (memGRK3ct-Rluc8)^19^ in the presence the S1 subunit of pertussis toxin (PTX) to prevent receptor-catalyzed nucleotide exchange. For wt G_i1_ we found that GDPβS-GTPγS exchange was much slower than GTPγS binding, consistent with the expectation that the rate of exchange is limited by GDPβS release^2^. In contrast, GDPβS-GTPγS exchange was much more rapid for G_i1_ 4A, and approached the rate of GTPγS binding (Fig. 2B). This observation suggests that G_i1_ 4A heterotrimers in cell membranes have a rapid rate of spontaneous GDPβS release, consistent with the idea that these mutants have structural and functional features of receptor- bound G proteins.

To test whether the properties of G_i1_ 4A are conserved across other G protein families, we constructed analogous Gα_s_ 4A, Gα_q_ 4A, Gα_12_ 4A subunits and examined interactions with cognate GPCRs for all four families; the α_2A_ adrenoreceptor (α_2_AR) for G_i1_ 4A, the β_2_ adrenoreceptor (β_2_AR) for G_s_ 4A, the M_3_ muscarinic acetylcholine receptor (M_3_R) for G_q_ 4A, and GPR35 for G_12_ 4A. We measured BRET between receptors and heterotrimers and directly compared wt Gα subunits with Gα 4A subunits. Responses to agonists and inverse agonists were studied in permeabilized cells in the presence of either apyrase or GDP. As expected, conventional allosteric coupling was observed for all four wt G proteins (Fig. 3A). Agonist- induced receptor-G protein association was modest in the presence of GDP, and much more robust in the absence of nucleotides. Because most GPCRs intrinsically favor inactive conformations inverse agonist responses mediated by wt G proteins were modest or undetectable, even in the absence of nucleotides (Fig. 3B). Unlike their wt counterparts, 4A heterotrimers spontaneously formed complexes with their cognate receptors, as indicated by high baseline BRET that was increased only slightly by agonists (Fig. 3A). Consistent with our results with M_4_R, BRET between G_i1_ 4A and α_2_AR was largely resistant to GDP, especially when agonist was present. BRET between the other 4A mutants and receptors we studied was completely resistant to GDP, suggesting that association of G_s_ 4A, G_q_ 4A and G_12_ 4A are completely decoupled from nucleotide exchange. BRET between receptors and 4A G proteins was partially sensitive to inverse agonists, consistent with disruption of active receptor-G protein 4A complexes that formed spontaneously. Notably, with the exception of rauwolscine in the presence of GDP, none of the inverse agonists tested was able to completely disrupt receptor-G protein 4A complexes (Fig. 3B).

**Figure 3.**
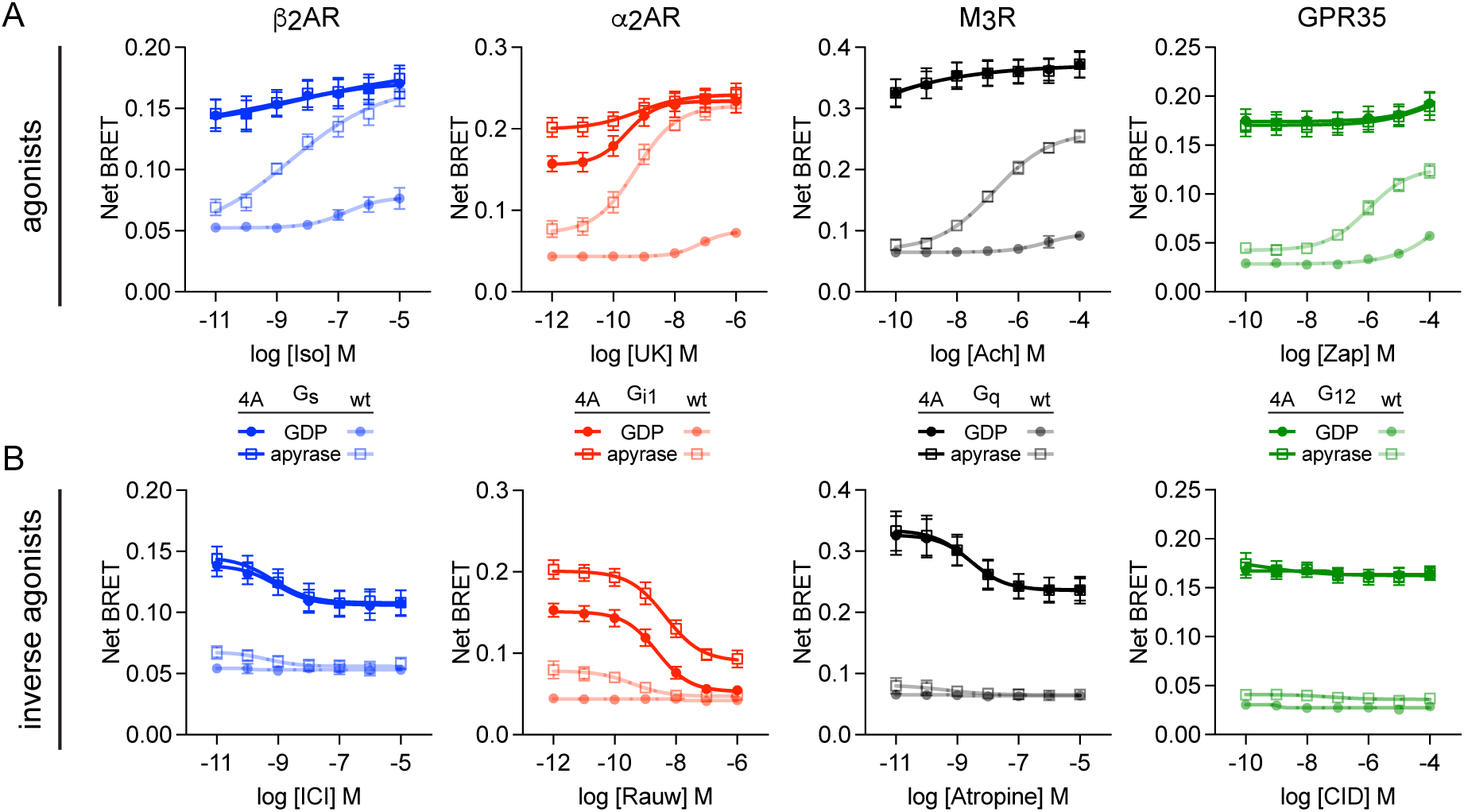
Nucleotide-decoupled G proteins bind spontaneously to cognate GPCRs. BRET between four representative Rluc8-tagged GPCRs and cognate wt and 4A G protein heterotrimers in response to agonists (*A*) and inverse agonists (*B*). Compared to wt G proteins, 4A mutants display high basal BRET that is only modestly enhanced by agonists and only partially reversed by inverse agonists. Receptors are β_2_ and α_2A_ adrenergic receptors (β_2_AR and α_2_AR), M_3_R acetylcholine receptors, and GPR35 (mean ± S.D.; *n*=5).

### Nucleotide-decoupled G proteins bypass a selectivity barrier for GPCR-G protein interaction

Taken together the foregoing results indicate that association of receptors and 4A mutant G proteins is decoupled from nucleotide exchange and, as a result of this decoupling, 4A G proteins associate spontaneously with unliganded cognate receptors. We reasoned that 4A mutants bypass the energetically-costly conformational changes associated with nucleotide release, and took advantage of this property to ask how such conformational changes contribute to receptor-G protein coupling selectivity. We hypothesized that if structural complementarity of the most stable intermediate during receptor-catalyzed nucleotide exchange (the nucleotide-free G protein-receptor complex) was the sole driver of selectivity, then nucleotide-decoupled G protein mutants would maintain their selectivity. In contrast, if earlier, less stable intermediates (nucleotide-bound G protein-receptor complexes) influence selectivity, then selectivity would be degraded with G proteins that essentially bypass these states.

To test this idea, we measured BRET between 32 ligand-activated GPCRs and six nucleotide- decoupled G protein subtypes (i1 4A, oA 4A, s 4A, q 4A, 12 4A, and 13 4A) in response to their respective agonists in intact cells. We found that basal net BRET was highest for cognate receptor-G protein 4A pairs for all receptors studied, demonstrating that selectivity between ligand-free receptors and G protein 4A subtypes remained partially intact. However, agonist- mediated activation of receptors showed that subtype selectivity was markedly degraded, such that the majority of receptors formed complexes with all six G protein 4A heterotrimers (Fig. 4; Fig. S2). As one example, M_4_R displayed robust agonist-induced BRET with 4A heterotrimers from all four G protein families, even though these receptors cannot activate wt G_s/olf_ and G_12/13_ heterotrimers and only weakly activate G_q_^20^. More stringent selectivity between 4A mutants was retained in a few receptors, and this subset included receptors that couple primarily to each of the four G protein families. For example, the G_s_-coupled bile acid receptor GPBAR exhibited more stringent selectivity for G_s_ 4A heterotrimers, associating with other G protein 4A mutants only in the presence of high concentrations of agonist. Notably, GPBAR is also highly selective for wt G_s/olf_ over other heterotrimers^20^. Likewise, 5-HT_2A_ serotonin receptors and GPR55 retained fairly strict selectivity for their cognate heterotrimers G_q_ 4A and G_13_ 4A, respectively. Interestingly, a few receptors selected one subtype yet rejected another within the same G protein family. For example, 5-HT_5A_ associated with G_i1_ 4A and G_q_ 4A but largely rejected all other subtypes including G_oA_ 4A, a subtype in the same family as G_i1_. Similarly, GPR55 preferentially formed complexes with G_13_ 4A over G_12_ 4A. In terms of agonist potency, we found that EC_50_ values for noncognate receptor-G protein 4A pairings were generally right-shifted in comparison to cognate pairings. Likewise, reversal of receptor-G protein 4A association after addition of inverse agonist or activation of a competing receptor was more rapid and complete for noncognate complexes (Fig. S3). These results indicate that activated receptors interact promiscuously with 4A mutant G protein heterotrimers. However, the extent to which selectivity is degraded varies widely between receptors, and a significant degree of selectivity is invariably retained.

**Figure 4.**
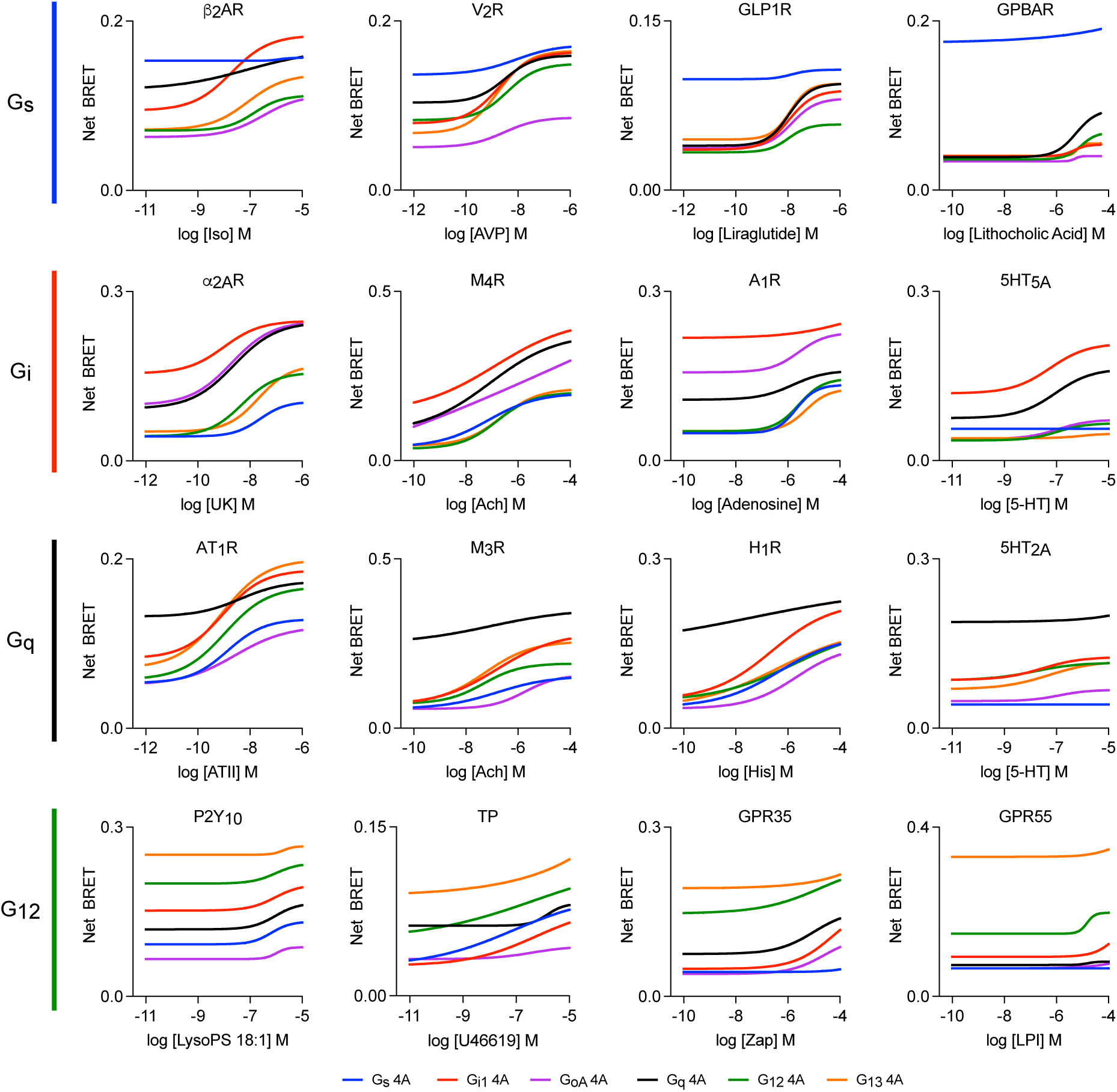
Nucleotide-decoupled G proteins bind promiscuously to noncognate GPCRs. BRET between sixteen representative Rluc8-tagged GPCRs and six different 4A G protein heterotrimers in response to agonists. Most receptors interact with all six 4A mutants in an agonist-dependent manner. Shown are least-square fits to a four-parameter concentration- response equation; data points have been removed for clarity. Concentration-response curves with data points from three independent experiments (± S.E.M.) are shown for all 32 receptors studied in Figure S2.

It was noteworthy that, except for 5-HT_5A_ and G_s_ 4A, all of the G_i/o_ coupled receptors that we studied formed agonist-dependent complexes with G_s_ 4A and G_q_ 4A heterotrimers (Fig. 4; Fig. S2). One mechanism that has been proposed to explain the selectivity of G_i/o_-coupled receptors is a limited displacement of transmembrane helix 6 (TM6) in the active state, which results in a binding pocket in the receptor TM core that is too small to accommodate the relatively bulky C termini of Gα_s_ and Gα_q_ subunits^21^. This hypothesis is consistent with the limited displacement of TM6 observed in many receptor-G_i/o_ complex structures^22-24^. However, from structural studies alone it is unclear if receptors with small TM6 displacements are unable to adopt larger displacements, or if instead larger displacements are possible but not captured when receptors are in complexes with G_i_ or G_o_. Our results showing robust association of G_i/o_-coupled receptors with G_s_ 4A and G_q_ 4A mutants are consistent with the latter explanation. To further support this conclusion, we verified that association of α_2_AR and M_4_R with G_s_ 4A required the Gα_s_ C terminus, and that neither receptor could detectably activate wt G_s_ heterotrimers (Fig. S4).

### Coupling selectivity of engineered Gα subunits

Gα subunits engineered to form stable complexes with active receptors have been valuable for solving structures of GPCR-G protein complexes. For example, variants of mini G proteins (mG) have been used to solve several GPCR-G protein complex structures by x-ray crystallography and cryo-EM^25^. These engineered Gα subunits lack the α helical domain altogether and incorporate several stabilizing mutations. One of the mutations in mG proteins, I^H5.4^ to A (Fig. 1), was found to uncouple guanine nucleotide binding and receptor association, such that GPCR- mG complexes would remain stable even in the presence of GDP or GTP ^26^. Because this property is shared with G protein 4A mutants we suspected that mG proteins might also have degraded selectivity for cognate GPCRs. We tested this idea by focusing on mini Gs, because most other mG proteins used for structural studies are chimeras that include significant sequence derived from Gα_s_. We constructed a variant of mini Gs (referred to here as midi Gs) with an intact αN helix, which provides a membrane anchor as well as the capacity to bind Gβγ. We also constructed an analogous subunit (midi Gs wtct) in which the uncoupling mutation was reverted to the wt sequence (A^H5.4^ to I). As was the case with G_s_ 4A, midi Gs (expressed together with Venus-Gβγ) interacted spontaneously with G_s_-coupled receptors such as the β_2_AR. This interaction was only slightly enhanced by agonist, was weakly diminished by inverse agonist, and was completely insensitive to the presence of GDP (Fig. 5A). In contrast, midi Gs wtct interacted with β_2_AR in an agonist- and GDP-sensitive manner, supporting the conclusion that the I^H5.4^ to A mutation decoupled nucleotide binding from receptor association^26^. We then tested the ability of midi Gs and midi Gs wtct to interact with noncognate receptors in intact cells. Similar to what we observed with G protein 4A mutants, midi Gs interacted well with noncognate receptors in an agonist-dependent manner (Fig. 5B). In contrast, midi Gs wtct interacted poorly with noncognate receptors. That two different mutations (4A insertion and I^H5.4^ to A) that decouple nucleotide binding from receptor association also degrade coupling selectivity strongly supports the notion that selectivity is partly determined by conformational changes that lead up to nucleotide release.

**Figure 5.**
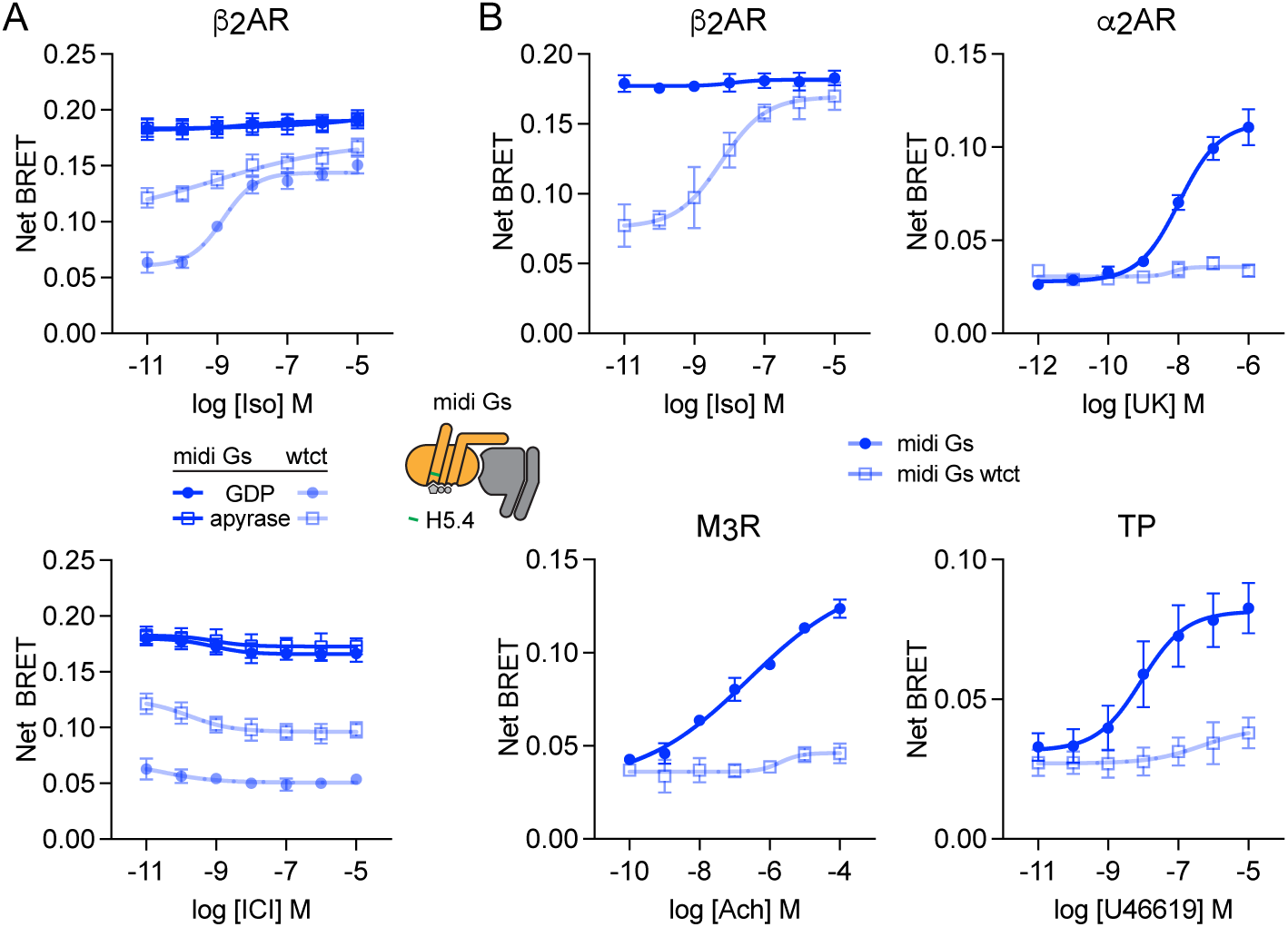
Nucleotide-decoupled midi Gs also loses selectivity. (*A*) In permeabilized cells midi Gs proteins bearing the nucleotide-decoupling mutation A^H5.4^ interact spontaneously with β_2_AR in a nucleotide-insensitive manner, whereas midi Gs bearing the wild-type residue in this position (wtct; I^H5.4^) interact with this receptor in a ligand- and nucleotide-sensitive manner (mean ± S.D.; *n*=5). (*B*) In intact cells midi Gs but not midi Gs wtct interacts promiscuously with G_i_-, G_q_- and G_12/13_-coupled receptors (α_2_AR, M_3_R and TP, respectively) in response to agonist stimulation (mean ± S.D.; *n*=3).

Dominant negative (DN) Gα subunits have also been valuable tools for structural determination of GPCR-G protein complexes^27^. DN G proteins have mutations that reduce nucleotide-binding affinity and enhance stability of ternary complexes. These functional properties prompted us to characterize their interaction with cognate and noncognate receptors, focusing on DN Gα_s_ and Gα_i1_. Compared to G protein 4A mutants and midi Gs, association of DN Gα_s_ and Gα_i1_ with cognate receptors (β_2_AR and α_2_AR) was more dependent upon agonist, and more sensitive to GDP, suggesting a milder nucleotide decoupling phenotype (Fig. S5A). At the same time, DN Gα_s_ and Gα_i1_ retained coupling selectivity to a greater degree than fully decoupled variants as receptors interacted with non-cognate DN G proteins only modestly, even in the absence of nucleotides (Fig. S5B). These results suggest that degradation of coupling selectivity across G protein variants correlates with the extent to which receptor association is decoupled from nucleotide binding.

### Constitutive conformational sampling is a near-universal feature of bona fide GPCRs

GPCRs are conformationally dynamic and can spontaneously transition between inactive and active states^28,29^. This property is likely to be responsible for constitutive G protein activation observed for some receptors. We observed a high basal BRET value (net BRET >0.1) for the cognate G protein 4A subtype for the 29 of the 32 receptors we studied (Fig. 6A), a sample that included receptors known to have low constitutive activity such as V_2_R (AVPR2) and α_2_AR (ADRA2A). This indicates that even receptors without appreciable constitutive activity sample active states to some extent, and suggests that nucleotide-decoupled G proteins are sensitive probes for detecting conformational sampling.

**Figure 6.**
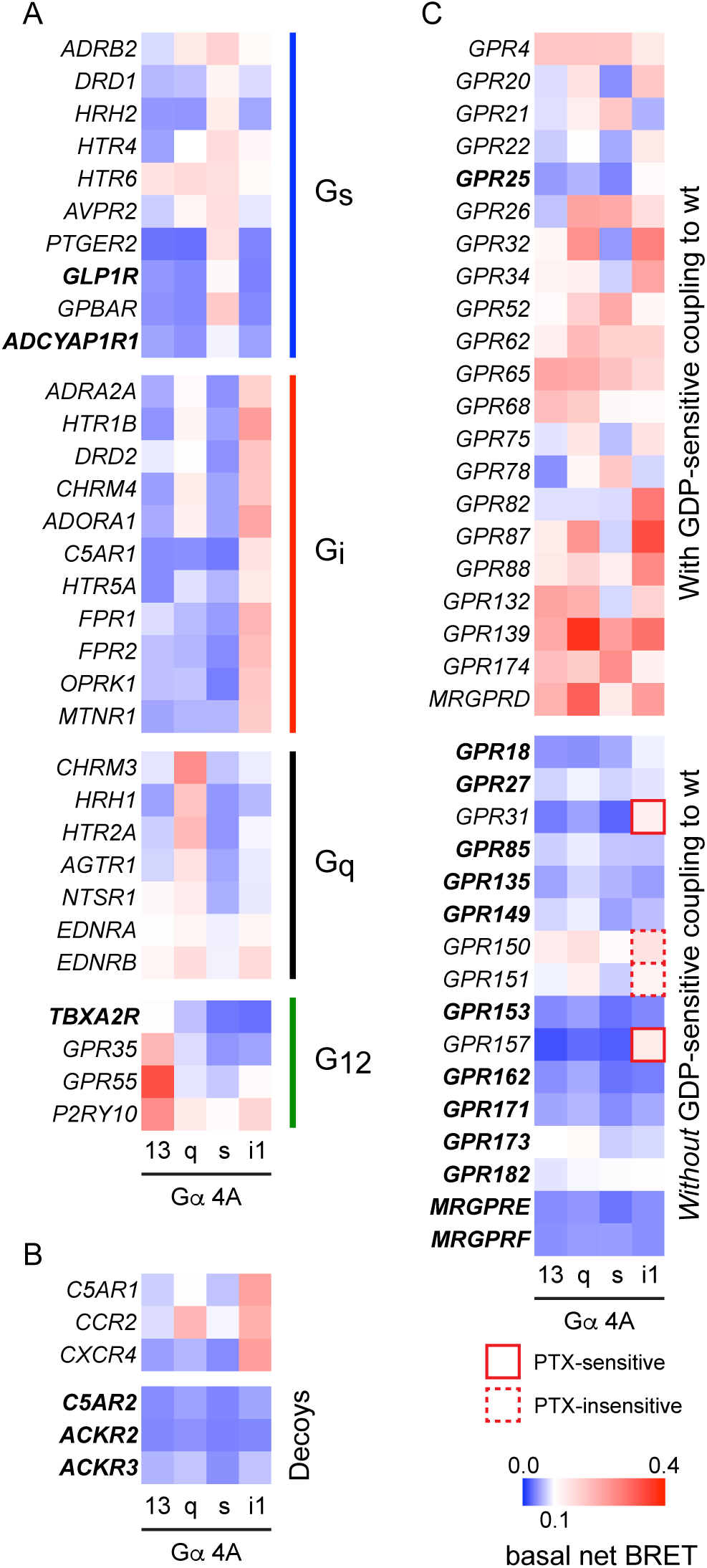
Many orphan GPCRs may be decoy receptors. (*A*) Heat maps showing basal BRET between 32 well-studied GPCRs and four different G protein 4A mutants. Only three of these receptors (GLP1R, ADCYAP1R1 and TBXA2, shown in ***bold italic*** font) show basal BRET that fails to exceed 0.1 with their cognate 4A mutant. (*B*) Three bona fide chemokine GPCRs show high basal BRET with G_i1_ 4A, whereas three known decoy receptors do not. (*C*) Most orphan receptors that show constitutive coupling to wt G proteins (*top*) also interact spontaneously with at least one G protein 4A mutant, whereas a significant fraction of orphan receptors that lack spontaneous coupling to wt G proteins also do not interact spontaneously with 4A mutants, suggesting that many of these receptors may be decoys. Exceptions were GPR150, GPR151, GPR157 and GPR31, all of which showed basal BRET that exceeded 0.1 with G_i1_ 4A, although this was sensitive to pertussis toxin (PTX) only for the latter two receptors. In all panels each cell represents the mean of 3 independent experiments.

We then asked if we could make use of this property to identify 7 transmembrane (7TM) proteins that are related to GPCRs but do not activate G proteins. Examples of this type of protein include so-called decoy receptors, which bind and sequester ligands yet fail to initiate G protein signaling. We predicted that such receptors would fail to interact with G protein 4A mutants. To test this prediction we first measured basal BRET between 4A mutants from each G protein family and three bona fide G_i/o_-coupled chemokine receptors (C5AR1, CCR2 and CXCR4), as well as three atypical chemokine receptors (C5AR2, ACKR2 and ACKR3) known to be decoys^30^. As expected, all three G_i/o_-coupled receptors showed high basal BRET to G_i1_ 4A heterotrimers. In contrast, none of the known decoy receptors showed significant basal interaction with any 4A mutant (Fig. 6B). We then applied this strategy to a panel of understudied orphan GPCRs (oGPCRs), many of which have not been shown to couple to any transducer. We divided this panel into two groups based on whether or not we previously detected conventional GDP-sensitive coupling to wt G proteins^31^. For oGPCRs shown previously to couple to wt G proteins 21 of 22 showed high basal BRET signals to at least one G protein 4A mutant, consistent with what we observed with liganded GPCRs. In contrast, of the oGPCRs without detectable GDP-sensitive coupling to wt G proteins, only 4 of 16 receptors (GPR31, GPR150, GPR151 and GPR157) showed basal net BRET that exceeded 0.1 with any 4A mutant (Fig. 6C). As all four of these receptors displayed high BRET to G_i1_ 4A we tested sensitivity of this signal to PTX, which nearly abolished basal BRET between G_i1_ 4A and GPR31 and GPR157, but did not significantly decrease energy transfer to the other two receptors (Fig. S5). These results suggest that GPR31 and GPR157 are both G_i_-coupled GPCRs, whereas GPR150 and GPR151 are unlikely to couple to G_i1_ but may couple to other G proteins. The twelve oGPCRs in this group that did not interact well with G protein 4A mutants may lack the ability to activate G proteins altogether, or may only be able to do so in the presence of an activating ligand.

## Discussion

The mechanism of selective coupling of GPCRs to G proteins has be studied for decades using a variety of experiment approaches, most of which have focused on the structural determinants of selectivity^3^. Most recently, solved structures of receptors bound to nucleotide-free G proteins have been used to predict residues in the receptor-G protein interface that contribute to selectivity. These studies have provided valuable insights into how individual receptors discriminate G protein subtypes, and collectively suggest that the conformation of the receptor-G protein complexes that are critical for selective coupling share many features with these empty-state complexes^5-8^. Nevertheless, it is recognized that structural studies are generally limited to stable complexes that exist only after GDP release, and that such complexes may not reflect the key intermediates that immediately precede nucleotide release. Therefore, it has been suggested that selective coupling may also be determined by transient intermediate states that are not amenable to standard structural methods^10,11^.

In the present study we considered the possibility that selectivity could be determined in part by the process whereby the receptor disrupts the hydrophobic core of the Gα subunit, concomitant with extension of the G protein C terminus. This conformational change is predicted to be both energetically costly ^15^ and critical for GDP release^14^. To test this idea we made use of G protein variants shown previously to bind to active GPCRs in the presence of guanine nucleotides^13^. In the case of G protein 4A mutants, the hydrophobic core of the Gα subunit is disrupted even though nucleotide is present. We reasoned that such mutants would effectively bypass some of the steps leading up to GDP release, and might therefore have less stringent coupling selectivity. Consistent with our hypothesis, we found that selectivity between GPCRs and nucleotide-decoupled G proteins was degraded to the point where most receptors could interact with G protein 4A subtypes from all four families.

Our results are consistent with a model of selectivity wherein the conformational changes in Gα necessary to promote GDP release represent a generic energetically-insurmountable barrier for noncognate GPCR-G protein pairs. Bypassing this barrier reveals a latent capacity for most GPCRs and G proteins to interact. One straightforward conclusion that can be drawn from this observation is that there are no absolute structural barriers to complex formation for the majority of potential GPCR-G protein pairs. At the same time, our results reinforce the importance of structural complementarity in GPCR-G protein coupling, as the conformational plasticity presumably conferred on the G protein by nucleotide-decoupling did not abolish selectivity. Indeed, some receptors (e.g. GPBAR) retained a high degree of selectivity between nucleotide- decoupled G proteins, consistent with highly variable selectivity mechanisms and stringency across GPCRs^32,33^. Our results do argue against the specific hypothesis that G_i/o_-coupled receptors generally reject G_s_ proteins because these receptors cannot accommodate the bulky C terminus of Gα_s_^21^. Most of the G_i/o_-coupled receptors that we studied bound to G_s_ 4A in the presence of agonist. Once again there were exceptions to this trend (e.g. 5-HT_5A_), further reinforcing the notion that different receptors have different selectivity mechanisms. In general, the observation that most receptors could interact to some extent with most G protein 4A subtypes suggests that selectivity is determined for many GPCR-G protein pairs by multiple weak interactions distributed across an extensive interface.

One surprising observation was that nucleotide-decoupled G proteins spontaneously formed complexes with cognate GPCRs without the need for agonist-induced activation. This finding suggests that most ligand-activated GPCRs constantly sample states other than the fully inactive state. This is not unexpected for receptors known to constitutively activate G proteins in the absence of an agonist, but our results suggest this property extends even to receptors that do not spontaneously activate G proteins. We made use of this property to probe a collection of orphan GPCRs and found that basal interaction with 4A mutants correlated well with GDP- sensitive coupling to wild-type G proteins. Most of the oGPCRs that did not couple detectably with wild-type G proteins also did not interact with 4A mutants. Two exceptions to this were GPR31 and GPR157, both of which showed PTX-sensitive interactions with G_i1_ 4A. These results agree with previous studies suggesting that GPR31 couples to G_i/o_ proteins^34^, but contrast with a report suggesting that GPR157 activates G_q/11_ proteins^35^. Almost all of the bona fide GPCRs that we studied showed robust constitutive signals with nucleotide-decoupled G proteins. Therefore, our results imply that many of the oGPCRs that did not interact with 4A mutants may in fact be analogous to decoy receptors, although we cannot rule out the possibility that ligand-induced activation of such receptors will ultimately be shown to promote G protein coupling.

In summary, nucleotide-decoupled G proteins bind promiscuously to agonist-activated GPCRs, but retain a preference for cognate receptors. To our knowledge this represents the first perturbation that degrades GPCR-G protein selectivity across receptors and G protein subtypes. Our findings are consistent with a model where coupling selectivity is determined at several steps during receptor-catalyzed nucleotide exchange, at several sites across an evolving receptor-G protein interface, and by mechanisms that differ widely between receptors.

## Materials and Methods

### Materials

Trypsin, culture media, PBS, DPBS, penicillin/streptomycin and L-glutamine were from GIBCO (ThermoFisher Scientific, Waltham, MA, USA). PEI MAX was purchased from Polysciences Inc. (Warrington, PA, USA). Digitonin, apyrase and GDP were purchased from MilliporeSigma (St. Louis, MO, USA). Coelenterazine h was purchased from Nanolight Technologies (Pinetop, AZ, USA) and furimazine (NanoGlo) was purchased from Promega (Madison, WI, USA).

### Plasmid DNA constructs

GPCR coding sequences were provided by Bryan Roth (University of North Carolina, Chapel Hill, NC; PRESTO-Tango Kit—#1000000068, Addgene, Watertown), MA, USA) ^36^, except for GPR139, which was a gift from Kirill Martemyanov^37^. For each receptor the coding sequence was amplified with a common forward primer (corresponding to a cleavable signal sequence) and custom reverse primer (corresponding to the receptor C terminus) and ligated into a pRluc8-N1 cloning vector. The design of memGRK3ct-Rluc8 was previously described^19^. Gα subunit plasmids were purchased from <u>cdna.org</u> (Bloomsburg University, Bloomsburg, PA. 4A insertion mutations were introduced to Gα subunits using the QuikChange Mutagenesis Kit (Agilent Technologies) and oligonucleotides (Integrated DNA Technologies) as primers. Gα_i1_ 4A was kindly provided by Heidi Hamm (Vanderbilt University, Nashville, TN). A plasmid encoding the Nluc-EPAC-VV cAMP sensor was kindly provided by Kirill Martemyanov (Scripps Research Institute, Jupiter, FL). All plasmid constructs were verified by Sanger sequencing. A plasmid encoding the S1 subunit of pertussis toxin (PTX-S1) was kindly provided by Stephen R. Ikeda (NIAAA, Rockville, MD, USA).

### Cell culture and transfection

HEK 293 cells (ATCC) were propagated in plastic flasks and on 6-well plates according to the supplier’s protocol. HEK 293 cells with targeted deletion of GNAS, GNAL, GNAQ, GNA11, GNA12, and *GNA13* (G protein three family knockouts; 3GKO) were derived, authenticated and propagated as previously described^17^. Cells were transfected in growth medium using linear polyethyleneimine MAX (PEI MAX; MW 40,000) at an nitrogen/phosphate ratio of 20 and were used for experiments 24-48 hours later. Up to 3.0 µg of plasmid DNA was transfected in each well of a 6-well plate.

### BRET assays

#### Measurement of coupling between receptors and G proteins

For G protein coupling in intact cells, HEK cells were transfected with a GPCR-Rluc8, Gα WT or 4A subunit, Venus-1-155-Gγ_2_, Venus-155-239-Gβ_1_, and pcDNA3.1(+) or PTX-S1 in a (1:10:5:5:5) ratio for a total of 2.6 µg of plasmid DNA in each well of a 6-well plate. After a 24- 48-h incubation, cells were washed twice in PBS and harvested by trituration in DPBS. Measurements were made from intact cells with either agonist, inverse agonist or DPBS control.

For G protein coupling in nucleotide-depleted cells, 3GKO cells were transfected with the same plasmids and ratio for the intact cell G protein coupling experiments. After a 48-h incubation, cells were washed twice with permeabilization buffer (KPS) containing 140 mM KCl, 10 mM NaCl, 1 mM MgCl_2_, 0.1 mM KEGTA, 20 mM NaHEPES (pH 7.2); harvested by trituration; permeabilized in KPS buffer containing 10 µg mL^−1^ high-purity digitonin; and transferred to opaque black 96-well plate. Measurements were made from permeabilized cells supplemented either with 100 µM GDP or 2U mL^−1^ apyrase, in both cases with either agonist, inverse agonist or KPS control.

#### GPCR competition assays

Cells were transfected with an untagged GPCR, GPCR-Rluc8, Gα 4A, Venus-1–155-Gγ_2_, and Venus-155–239-Gβ_1_ in a (3:3:8:4:4) ratio for a total of 2.2 µg of plasmid DNA in each well of a 6- well plate. After a 48-h incubation, cells were washed twice with DPBS, harvested by trituration, and transferred to opaque white 96-well plates.

#### Nluc-EPAC-VV cAMP assay

For testing G_s_ activation by G_i_-coupled receptors (Fig. S4), 3GKO cells were transfected with a GPCR, Gα_s_ subunit, Gβ_1_, Gγ_2_, Nluc-EPAC-VV, pcDNA3.1(+), and PTX-S1 in a (15:5:10:10:1:49:10) ratio. After a 24-h incubation, cells were washed twice with DPBS, harvested by trituration, and transferred to opaque black 96-well plates.

#### Nucleotide exchange rate assay

Cells were transfected with a memGRK3ct-Rluc8, Gα 4A, Venus-1–155-Gγ_2_, Venus-155–239- Gβ_1_, and PTX-S1 in a (3:10:5:5:3) ratio for a total of 2.6 µg of plasmid DNA in each well of a 6- well plate. After a 48-h incubation, cells were washed twice with KPS and harvested by trituration in KPS containing digitonin. Permeabilized cells were supplemented with 2U mL^−1^ apyrase with or without 10 µM GDPβS for at least 30 minutes. The rate at which free Gβγ- Venus is released upon addition of 10 µM GTPγS in the absence of GDPβS reflects GTPγS incorporation into nucleotide-free Gα, whereas in the presence of GDPβS the rate of GTPγS incorporation and Gβγ-Venus release is limited by GDPβS dissociation (nucleotide exchange).

### BRET measurements

Steady-state BRET and luminescence measurements were made using a Mithras LB940 photon-counting plate reader (Berthold Technologies GmbH). Kinetic BRET and luminescence time course measurements were made using a Polarstar Optima plate reader (BMG Labtech). Coelenterazine h (5 µM) or furimazine (1:1,000) were added to all wells immediately prior to making measurements with Rluc8 and Nluc, respectively. Raw BRET signals were calculated as the emission intensity at 520–545 nm divided by the emission intensity at 475–495 nm. Net BRET signals were calculated as the raw BRET signal minus the raw BRET signal measured from cells expressing only the Rluc8 donor.

## Acknowledgments

We thank Aska Inoue for providing CRISPR-modified cells lacking Gα subunits, and Heidi Hamm, Steve Ikeda, Kirill Martemyanov and Bryan Roth for providing plasmid DNA. This study was supported by NIH grants GM130142 and GM145284 (N.A.L.), and a PhRMA Foundation Predoctoral Fellowship in Drug Discovery (W.J.).

## Supplementary Figures

**Figure S1.**
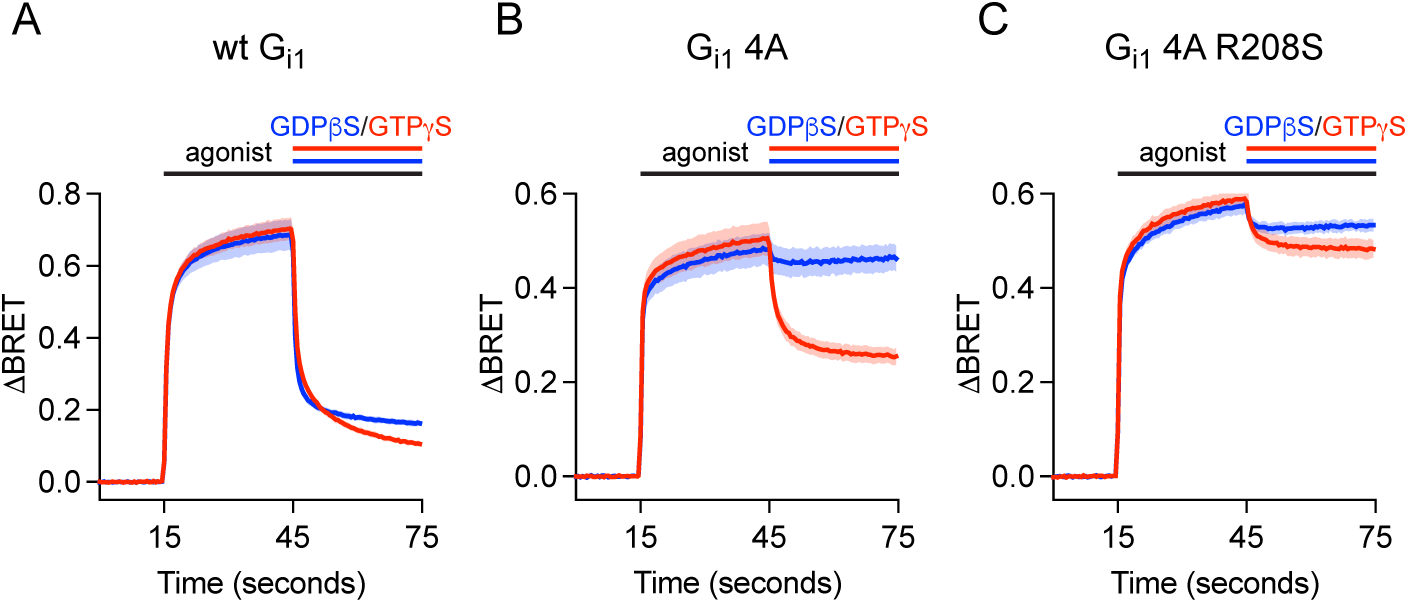
G_i1_ 4A interaction with the ligand-activated M4R is decoupled from nucleotide binding. *A)* BRET between M_4_R-Rluc8 and wild-type (wt) G_i1_ heterotrimers induced by the agonist acetylcholine (100 µM) in the absence of nucleotides was largely reversed by the addition of GTPγS or GDPβS (mean ± 95% C.I.; *n*=12). *B)* The response for G_i1_ Ins4A was insensitive to GDPβS and partially sensitive to GTPγS (mean ± 95% C.I.; *n*=16-20). *C)* βγ release-deficient G_i1_ Ins4A R208S nearly abolished GTPγS sensitivity of the receptor-G protein complex, suggesting that the nucleotide sensitivity of G_i1_ Ins4A is at the level of heterotrimer dissociation, not the allostery between receptor- and nucleotide-binding interface (mean ± 95% C.I.; *n*=12).

**Figure S2.**
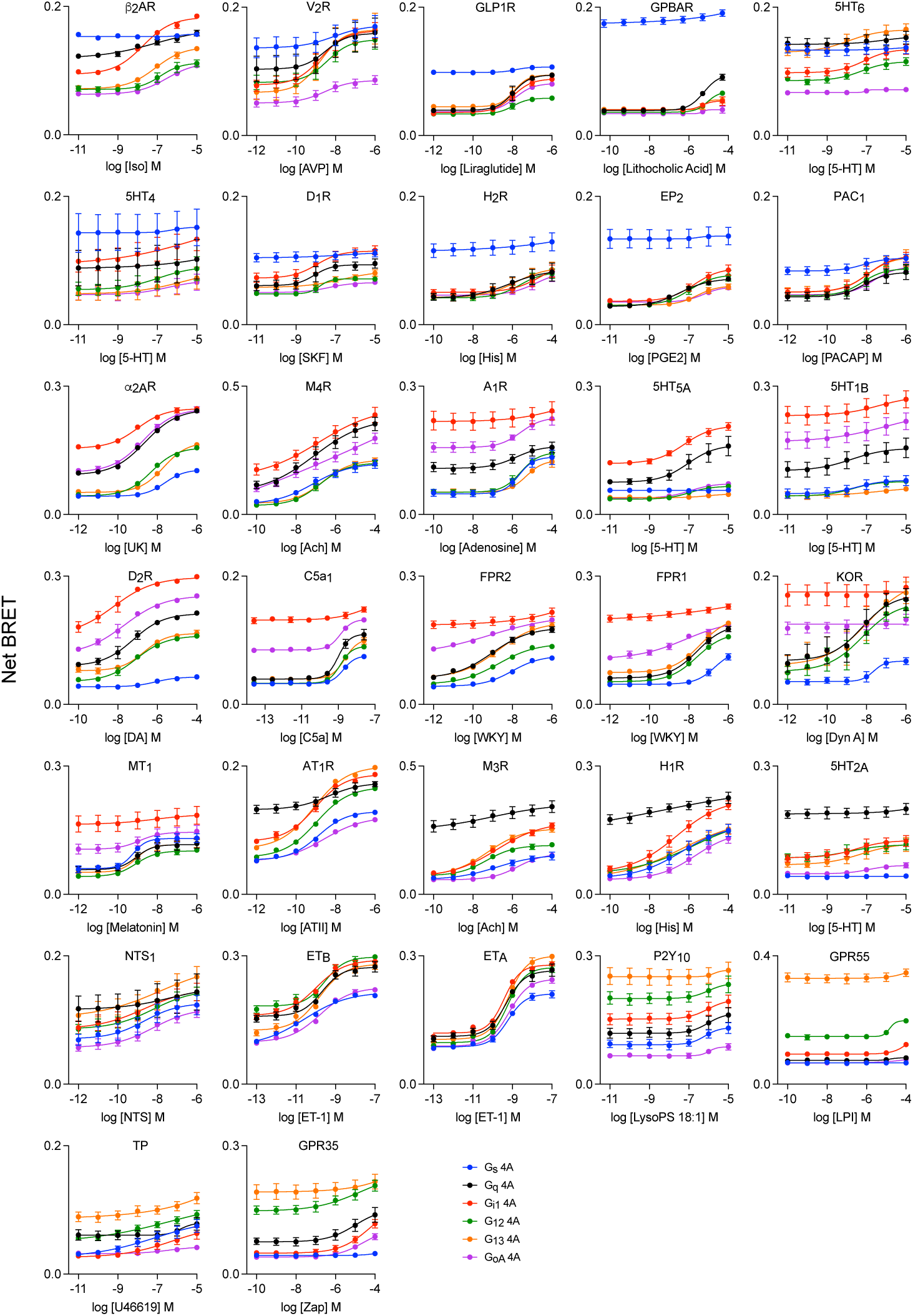
Concentration-response curves of BRET between six different 4A G protein heterotrimers and all 32 studied Rluc8-tagged GPCRs in response to agonists with data points (mean ± S.E.M.; *n*=3). Receptors are arranged according to their putative primary G protein coupling families.

**Figure S3.**
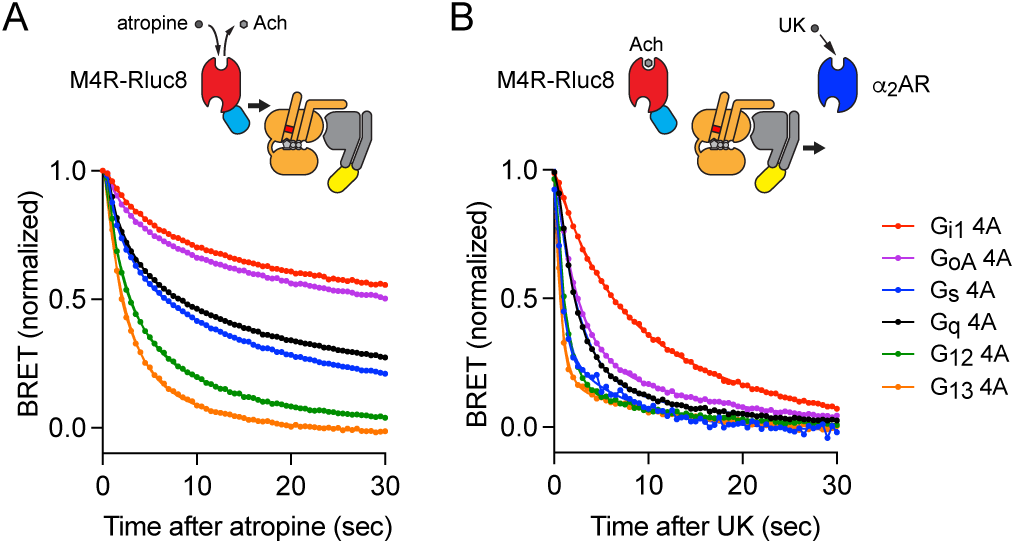
Noncognate receptor-G protein 4A complexes are more readily disrupted than their cognate counterpart. *A)* The time course of BRET in intact cells between M4R-Rluc8 preactivated by acetylcholine (100 µM) and Gα 4A βγ-Venus in response to 10 µM Atropine reveals slower decrease for G_i/o_ 4A heterotrimers than noncognate counterparts. Traces represent change of BRET normalized to baseline and are the average of 12 replicates from 3 independent experiments. *B)* Likewise, the time course of BRET in intact cells between M4R-Rluc8 preactivated by acetylcholine (100 µM) and Gα 4A βγ-Venus in response to activation of α_2_AR by its agonist UK-14,304 (1 µM) is slower for cognate interaction than noncognate interactions. Traces represent change of BRET normalized to baseline and are the average of 5-9 replicates from 3 independent experiments.

**Figure S4.**
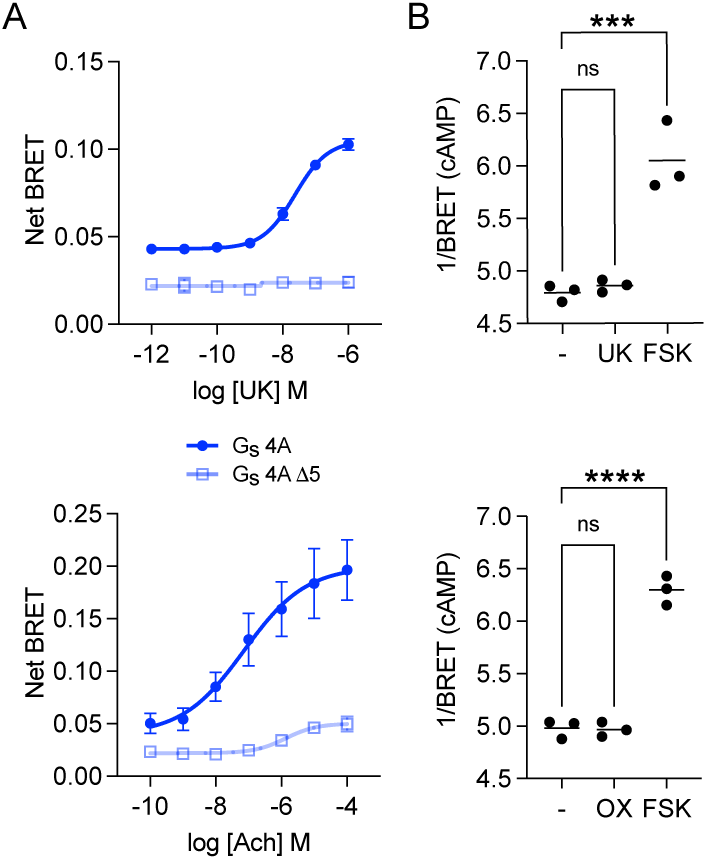
G_i_-coupled receptors accommodate C terminus of nucleotide-decoupled G_s_ heterotrimers yet cannot activate wt G_s_. *A)* Agonist-induced increase in BRET between Rluc8-tagged G_i_-coupled receptors (α_2_AR (*top*) and M_4_R (*bottom*)) and Gα 4A βγ-Venus are largely occluded when Gα_s_ 4A is truncated (mean ± S.D. n=3-5). *B)* Upon agonist-induced activation, α_2_AR (*top*) and M_4_R (*bottom*) cannot activate wt G_s_ heterotrimers to increase cAMP as Forskolin can (n=3). P < 0.05, one-way ANOVA (Dunnett’s test).

**Figure S5.**
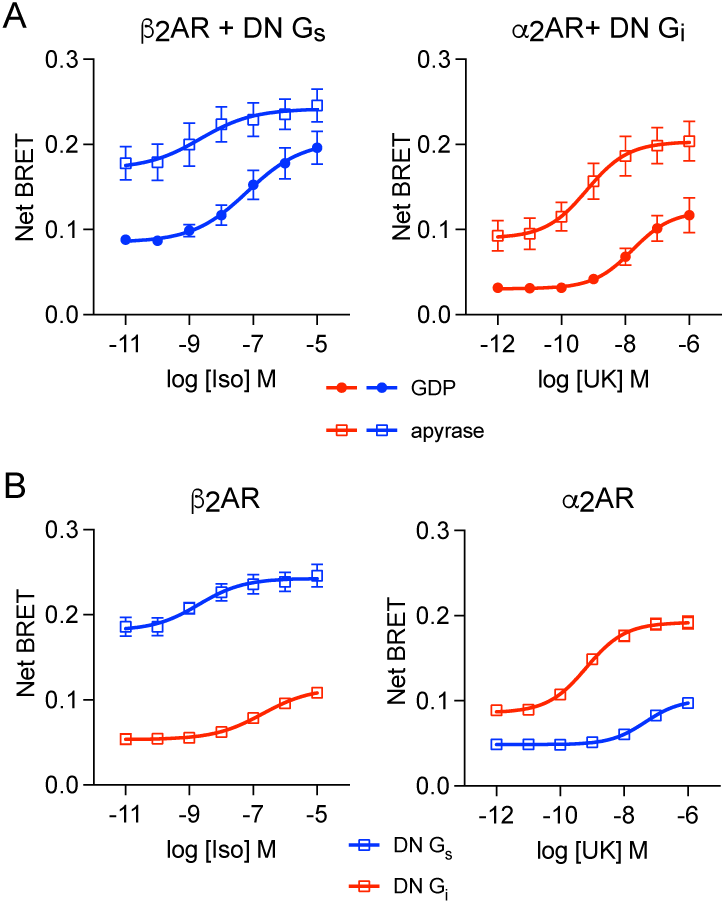
Dominant negative (DN) G proteins have a milder nucleotide decoupling phenotype. *A)* In permeabilized cells association between DN Gα_s_ and Gα_i1_ with their cognate receptors (β_2_AR and β_2_AR) are both agonist- and nucleotide-dependent albeit to a lesser degree in comparison with G 4A mutants and midi Gs (mean ± S.D. n=5). B) DN Gα_s_ and Gα_i1_ associated with noncognate receptors modestly in the absence of nucleotides (mean ± S.D. n=4).

